# Dynamic changes in rhythmic and arrhythmic neural signatures in the subthalamic nucleus induced by anaesthesia and intubation

**DOI:** 10.1101/762260

**Authors:** Yongzhi Huang, Kejia Hu, Alexander L. Green, Xin Ma, Martin J. Gillies, Shouyan Wang, James J. Fitzgerald, Yixin Pan, Sean Martin, Peng Huang, Shikun Zhan, Dianyou Li, Huiling Tan, Tipu Z. Aziz, Bomin Sun

**Affiliations:** Nuffield Department of Surgical Sciences, John Radcliffe Hospital, University of Oxford, Oxford OX3 9DU, UK; Center of Functional Neurosurgery, Ruijin Hospital, Shanghai Jiao Tong University School of Medicine, Shanghai 200025, China; Department of Anesthesiology, Ruijin Hospital, Shanghai Jiao Tong University School of Medicine, Shanghai 200025, China; Institute of Science and Technology for Brain-Inspired Intelligence, Fudan University, Shanghai 200433, China; Medical Research Council Brain Network Dynamics Unit at the University of Oxford, Oxford OX1 3TH, UK; Nuffield Department of Clinical Neurosciences, John Radcliffe Hospital, University of Oxford, Oxford OX3 9DU, UK

**Keywords:** propofol, anaesthesia, subthalamic nucleus, power law, local field potential

## Abstract

**Background:** Subcortical structures including the basal ganglia have been proposed to be crucial for arousal, consciousness, and behavioural responsiveness. However, how basal ganglia contributes to the loss and recovery of consciousness during anaesthesia has not been well characterized.

**Methods:** In this study, using local field potentials (LFPs) from subthalamic nucleus (STN) and scalp electroencephalogram in 12 Parkinson’s disease patients, we investigate STN neural signatures during propofol general anaesthesia and during intubation as an arousal intervention in anaesthesia.

**Results:** Propofol-induced anaesthesia resulted in changes in multiple frequency bands in STN LFPs, including increased low-frequency activities (slow-wave oscillation, delta, theta, and alpha bands) and decreased higher-frequency activities. This was also accompanied by increased STN-frontal cortical coherence in alpha frequency band. Beta and high-gamma activities in the STN temporally increased during intubation compared to the status of loss of consciousness. We also show that the dynamic changes in the high frequency activities (80-180 Hz) in STN LFPs induced by propofol and intubation correlated with power-law exponent in the power spectra between 2 and 80 Hz.

**Conclusions:** Our findings suggest that anaesthesia and intubation induced changes in the STN LFPs in multiple frequency bands. They are also consistent with the hypothesis that the power-law exponent in the power spectra between 2 and 80 Hz reflect the excitation/inhibition balance in the STN, which is modulated by anaesthesia and intubation, and further modulate the high frequency activity.

## Introduction

Understanding of the alteration of brain states under general anaesthesia has been significantly advanced by recent electrophysiology and functional neuroimaging studies. Anaesthetic-induced unconsciousness is thought to be associated with changes in different neural oscillations in cortical brain areas, commonly monitored by scalp EEG. For example increase in global slow-wave oscillation (SWO) power ^1, 2^, increase in frontal alpha power ^1, 3, 4^, burst suppression ^5, 6^, and changes in coherence ^1, 3^ were observed during induction of unconsciousness with propofol, one of the most commonly used aesthetic drugs. Numerous neuroimaging studies have also shown that several subcortical brain structures are involved in altered arousal and unconsciousness ^7–9^, and it is suggested that the disruption of communication between the cortical and subcortical structures play a crucial role in anaesthetic-induced unconsciousness^8, 10, 11^. Therefore, investigating subcortical changes and neuronal network-level effects induced by anaesthetic drugs has become compelling and necessary for better understanding the neuronal mechanism of how general anaesthetics cause loss of consciousness ^12–14^.

The basal ganglia, composed of striatum, globus pallidus, subthalamic nucleus (STN) and substantia nigra, are generally involved in a wide range of motor and cognitive processes that operate on the basis of wakefulness. A growing amount of studies start to highlight the importance of cortico-basal ganglia-thalamo-cortical network in arousal and consciousness ^15–18^. Recent investigation on the connectivity within the cortico-basal ganglia-thalamo-cortical network has shown that propofol-induced unconsciousness modulated the pallidal connectivity to the posterior cingulate cortex, suggesting the role of pallido-cortical connectivity in altered states of consciousness ^17^. Recent study also showed that propofol significantly modulated the oscillatory neuronal activity in the beta frequency band in the STN ^19^. However, electrophysiological brain activity consists of rhythmic oscillation and arrhythmic activity indicative of the oscillatory and fractal dynamics, respectively, arising from likely distinct neural basis ^20–22^. Up to now, most of the electrophysiological research on anaesthesia has focused on narrow-band oscillatory activity patterns. Meanwhile, arrhythmic brain activity coexists with brain oscillations and accounts for the majority of the signal power recorded in LFP/EEG/MEG, but our understanding of it remains very limited. The arrhythmic activity often appears with two spectral profiles: broad band and scale free ^23^. The power spectrum of scale-free dynamics tends to follow a power-law distribution: P~1/f^*β* ^20, 22, 24, 25^, which means the power (P) tends to fall off with increasing frequency (f) with *β* the power-law exponent. Increasing evidence has emerged suggesting that the scale-free features can uncover the functional involvement of neuronal activity in specific states. The power-low exponent has been shown to be modulated by sleep ^26^, neurological and psychiatric disorders ^27, 28^. The changes of power-low exponent during propofol-induced anaesthesia were also observed from human EEG and macaque electrocorticography (ECoG) ^29, 30^.

Here, we recorded electrophysiological signals from STN and frontal cortex simultaneously in patients with Parkinson’s disease (PD) undergoing deep brain stimulation (DBS) surgery. We compared the neuronal signatures including oscillatory activity, power-law distribution, functional connectivity in these brain areas under awake, propofol-induced unconsciousness, and under tracheal intubation in anaesthesia. We hypothesize that both oscillatory and scale-free arrhythmic activities in the STN, and the functional connectivity with cortex will be modulated by altered consciousness in general anaesthesia. In particular, we hypothesize that the temporal dynamics of the power-law exponent in the STN local field potential (LFP), which is hypothesized to reflect the excitation/inhibition balance, will also reflect the altered consciousness and arousal.

## Methods

### Subjects

Twelve patients with advanced PD (24 hemispheres, 3 females) undergoing bilateral DBS surgery of the STN were included in this study. All subjects gave their written informed consent and the local ethics committee approved the study. Details of the patients are reported in Table 1. All recordings in this study were performed when the patients underwent the second stage operation to implant with a subcutaneous pulse generator under general anaesthesia.

**Table 1.** Demographics and relevant clinical information

| Subject | Surgical centre | Age/Sex | Weight/kg | Medication before recordings (LDED) | Channel selection |
| --- | --- | --- | --- | --- | --- |
| 1 | Oxford | 43/F | 90 | 900 | L2-3;R1-2 |
| 2 | Oxford | 57/F | 101 | 1200 | L0-1;R0-1 |
| 3 | Oxford | 44/M | 71 | 775 | L0-1;R1-2 |
| 4 | Oxford | 65/M | 79 | 1300 | L1-2;R1-2 |
| 5 | Shanghai | 56/M | 71 | 1600 | L0-1;R1-2 |
| 6 | Shanghai | 64/M | 65 | 900 | L1-2;R1-2 |
| 7 | Shanghai | 73/M | 65 | 750 | L1-2;R1-2 |
| 8 | Shanghai | 65/F | 67 | 937.5 | L0-1;R0-1 |
| 9 | Shanghai | 57/M | 80 | 963 | L1-2;R1-2 |
| 10 | Shanghai | 57/M | 78 | 1027.5 | L1-2;R1-2 |
| 11 | Shanghai | 46/M | 63 | 1250 | L1-2;R1-2 |
| 12 | Shanghai | 34/M | 68 | 600 | L1-2;R1-2 |

LDED: L-DOPA daily equivalent dose.

### Anaesthetic procedures and data acquisition

The technique for induction of anaesthesia was at the discretion of the attending anaesthetist. After establishment of standard monitoring (electrocardiogram, noninvasive blood pressure, and pulse oximeter), all patients received sufentanil (0.5 ug kg^−1^) 60 s before propofol. Loss of consciousness (LOC) was achieved by i.v. propofol (1 – 2 mg kg^−1^). Time to LOC was determined by checking every 5 s for the loss of verbal response and eyelash reflex. After LOC, rocuronium (0.6 mg kg^−1^) was given to patients 142 s ± 51 s (range 80 s – 258 s) before tracheal intubation.

Electrophysiological signals from bilateral STN and frontal cortex were obtained from 3 min before the start of anaesthetic procedure to 2 min after intubation (11.6 ± 2.7 mins in total for each patient). Signal acquisition was carried out using TMSi Refa system (TMS International, Netherlands) with a sampling rate of 2048 Hz (Oxford) or a home-built device with a sampling rate of 1000 Hz (Shanghai). LFP signals were recorded with the four contacts of the DBS electrode (0, 1, 2, 3: 0 being the deepest contact) using monopolar configuration (Oxford) or bipolar configuration (0-1, 1-2, and 2-3) (Shanghai), with a reference electrode placed on the surface of the mastoid. Bipolar frontal EEG (FP_1, or 2_ – Fz) was recorded according to the international 10-20 EEG system.

### Data pre-processing

Bipolar LFPs (Oxford) were subsequently derived offline by subtracting the monopolar recordings between neighbouring contacts on each electrode. LFP and EEG data were resampled at 1000 Hz, high-pass filtered at 0.2 Hz, and notch-filtered at 50 Hz and the harmonics. The recordings were visually inspected and those data with noise and artefacts including excessive movement, muscle artefacts, and jump artefacts, were excluded from further analysis. This led to 4 out of the 24 STN LFPs during intubation and 4 out of the of 12 frontal EEG to be removed from final analysis. The selection of bipolar channel (see details in Table 1) for further LFP analysis was based on the review of pre- and post-operative imaging and post-operative clinical programming by the surgical team. Forty second of continuous data from each of the following three states were considered for each subject: (1) awake state, before anaesthetic induction, (2) loss of consciousness (LOC) state, after loss of eyelash reflex and before intubation, and (3) intubation state, beginning at 10 s after starting intubation.

### Spectral analysis

Power spectra of the electrophysiological signals (bipolar LFPs and EEGs) were calculated for all subjects in each state (awake, LOC, intubation) with the multitaper method using the Chronux toolbox ^31^. Multitaper parameters were set using time window lengths of T = 2 s, time-bandwidth product TW = 2, and number of tapers K = 3, resulting in a spectral resolution of 2 Hz when quantifying the average power spectra of the signal from different consciousness states. Time-frequency decomposition was also estimated to quantify the temporal dynamics of the power spectra by using the multitaper spectrogram method, with window lengths of 2 s and overlap of 1.5 s, TW = 2, and K = 3.

### Coherence analysis

To describe the connectivity between STN and frontal cortex, the magnitude squared coherence between STN LFPs and frontal EEG was estimated using the multitaper method, with window lengths of 4 s, TW = 4, and K = 7. The magnitude squared coherence estimate is the normalized cross-spectral density by the power spectral density with values between 0 and 1 that indicates the degree of covariability between the two signals over different frequency ranges. Imaginary coherence, which is only sensitive to synchronization of two signals that are time-lagged to each other, was further estimated to remove volume conduction of common signals. We derived the imaginary coherence between STN LFPs and frontal EEG using similar parameters used for coherence analysis.

### Power-law scaling analysis

Arrhythmic activities in the brain local field potentials tend to exhibit a 1/f-like spectrum: P~1/f^*β*. To characterize the property of these neural signals, the power spectrum of neural signals is often fitted by a linear function in a log-log coordinate space, where the slope (*β*) provides an estimate of the underlying power-law (or frequency-scaling) exponent. Here we investigated how the power-low exponent in STN LFP and frontal cortical EEG are modulated by alteration of conscious states under general anaesthesia. In addition, rhythmic brain oscillations appear as local peaks that rise above the power-law distribution in the power spectrum, which can confound the real scale-free dynamics, highlighting the importance of making a careful dissociation between rhythmic oscillation and arrhythmic activity ^20, 25, 32^. To estimate an accurate power-law exponent independent from any narrow band oscillations, we first adopted the irregular-resampling auto-spectral analysis (IRASA) method, to separate the fractal from oscillatory components of the power spectrum ^32^. This method takes advantage of the self-affinity property of a fractal (i.e. scale-free) time series, i.e., the statistical distribution of the data remains unchanged when resampled at different scales. Moreover, to further mitigate the potential effect of remaining spectral peak unsuccessfully removed by the IRASA method, the fractal power spectrum across a frequency range, excluding a narrower frequency band with local spectral peaks, was fitted. Then, the power-law exponent is estimated from the slope of the linear fit.

Here, a sliding time window with a length of 2 s and overlap of 1 s was applied for STN LFP and frontal EEG. Within each time window, the fractal power spectrum and corresponding power-law exponent was estimated. We further averaged the time courses of the power-law exponents within distinct behavioural states and compared the results across states.

### Statistical analysis

For STN LFPs, we determined whether or not there were significant differences of power in oscillatory activities in each frequency band and arrhythmic power-law exponent between awake and LOC states, and between LOC and intubation states. Before comparison, raw values of power and power-law exponent across subjects in each state were examined for deviations from normality using the Kolmogorov-Smirnov test. The power and exponent values were compared between states using the Wilcoxon signed ranks test if the data is not normally distributed or paired t-test if it is normally distributed. For frontal EEG, statistical significance of the differences of power and power-law exponent between LOC and intubation states were assessed. The imaginary coherence between STN LFPs and frontal EEG in LOC and intubation states was also compared. Resulting p-values were accordingly corrected for multiple comparisons with the false discovery rate (FDR) procedure across bands and states. All the signal processing and statistical analysis were conducted using MATLAB (Version 9.1, MathWorks, Inc., Natick, MA, USA).

## Results

### Increased low frequency oscillatory activity and reduced high frequency activity in the STN during anaesthesia compared to awake

We first compared the spectrograms from STN LFP and frontal EEG signals during the transition from awake to unconsciousness. The spectrograms showed dynamic spectral changes during induction of unconsciousness with the most dominant characteristics being the increased power in the alpha oscillation (8-14 Hz) and slow wave oscillation (SWO, 0.2-1.5 Hz) in both STN and frontal cortex at loss of consciousness (LOC) (Fig. 1A). In addition, there was also significant decrease in power in the high beta (22-40 Hz), and gamma (40-80 Hz) frequency bands in the STN at LOC (Fig. 1A). Group-level spectra (n = 24, twelve subjects) revealed that, compared with the awake state, the power in low frequencies including SWO, delta (1.5-4 Hz), theta (4-8 Hz), and alpha oscillations significantly increased, meanwhile, the power in higher frequencies including high-beta, gamma, high-gamma (80-180 Hz) bands significantly decreased in the STN at LOC state (SWO: *p* = 0.0288, delta: *p* = 0.0017, theta: *p* = 0.0006; alpha: *p* = 0.0005; high-beta: *p* = 0.0021, gamma: *p* = 0.0032, high-gamma: *p* = 0.0017; FDR correction) (Fig. 1B and C). In addition, we also noticed that during the awake state, frontal EEG showed broadband elevated high-frequency activities (Fig. 1C) which could be induced by muscle or eye movement ^33, 34^. Therefore, we excluded power and power-law analysis for awake EEG in further analysis.

**Figure 1.**
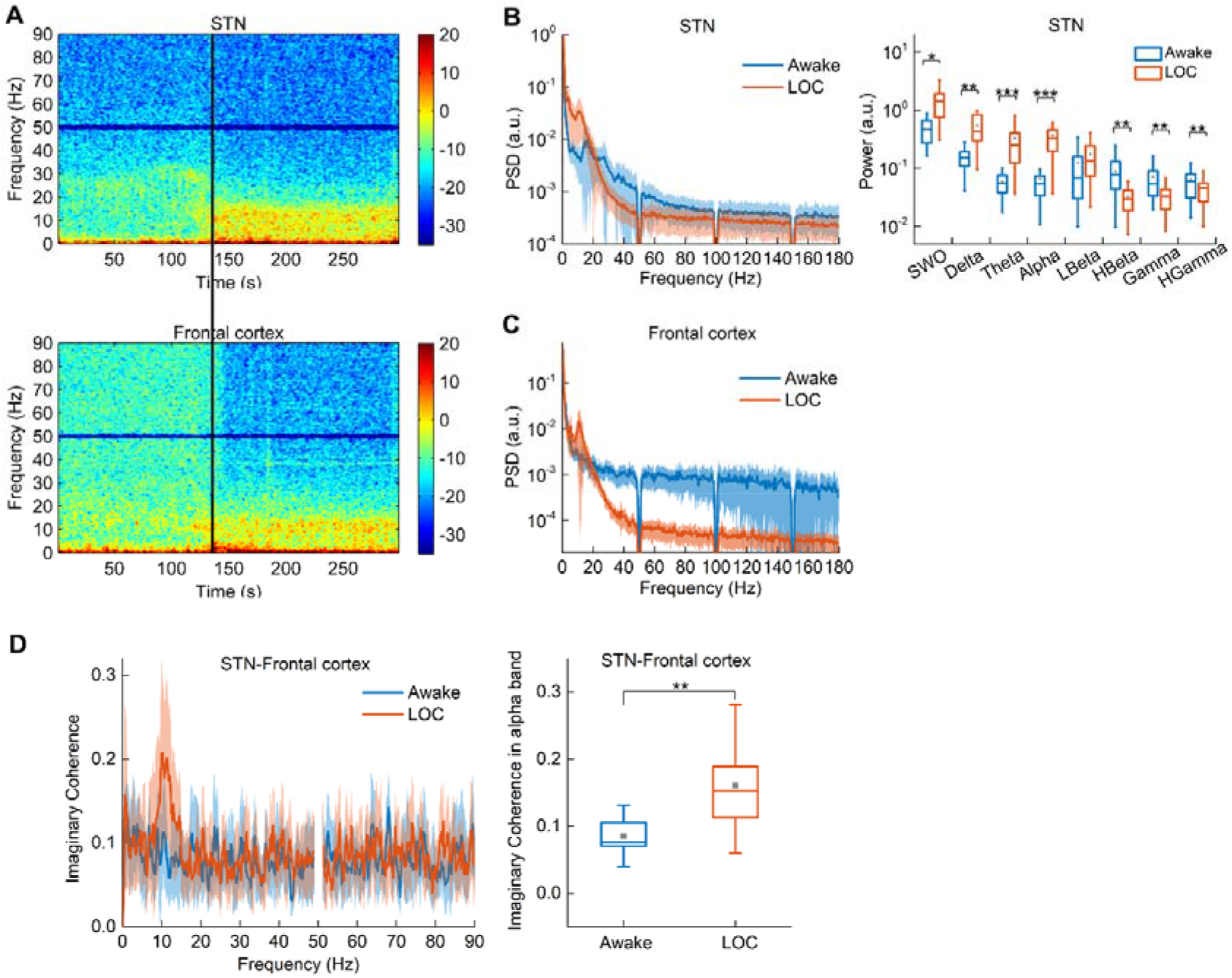
Increased low frequency activity and reduced high frequency activities in STN during LOC compared to awake, with the alpha band activity coherent with the frontal cortex. (A) Representative time-frequency spectrograms of STN LFPs and frontal cortical EEG during the induction of unconsciousness. The vertical black line marks the time of loss of eyelash reflex indicating LOC. (B) Group spectra and boxplot showing power in different frequency bands of STN LFPs across all subjects in awake and LOC states. (C) Group spectra of frontal EEG across subjects in awake and LOC states. (D) Group imaginary coherence between STN and frontal cortex across subjects in awake and LOC states. *, p<0.05; **, p<0.01; ***, p<0.001.

We next investigated how propofol affects subcortical-cortical communication by evaluating the coherence between STN LFPs and frontal cortical EEG. Group-level imaginary coherence analysis (n = 16, eight subjects) showed that at LOC, there was a significant increase in STN-frontal cortex coherence but only in the alpha range (8-13 Hz) (*p* = 0.0049) (Fig. 1D).

### Temporal dynamics of neural activity in STN measured by power-law distribution is differential in awake and LOC states

The power spectra of the STN LFPs, plotted in a log-log coordinate space (Fig. 2A), roughly followed a straight descending line, with local peaks corresponding to well-known neural oscillations rising above this line. Figure 2A also shows the fractal components of the power spectra (after removing the oscillatory components) for an example LFP signal from STN at awake and LOC states. However, as broadband spectral peaks could not be removed completely, the remaining fractal components of the power spectrum still had a “hump” in some frequency ranges, although reducing oscillatory activity. Therefore, we focused on the linear fitting of the fractal component of power spectrum across a frequency range (2–80 Hz), excluding 8–40 Hz range which covers most oscillatory activity (e.g. STN beta oscillation at awake and alpha oscillation at LOC). Group-level power-law exponents across all subjects (n = 24, twelve subjects) estimated from the slope of the linear fitting for each tested STN showed that, averaged exponent value in the STN significantly increased from 1.12±0.27 (range 0.65-1.93) in awake state to 1.83±0.29 (range 1.26-2.22) in LOC state (*p* < 0.0001) (Fig. 2B), indicating steeper reduction of power with increasing frequency in LOC.

**Figure 2.**
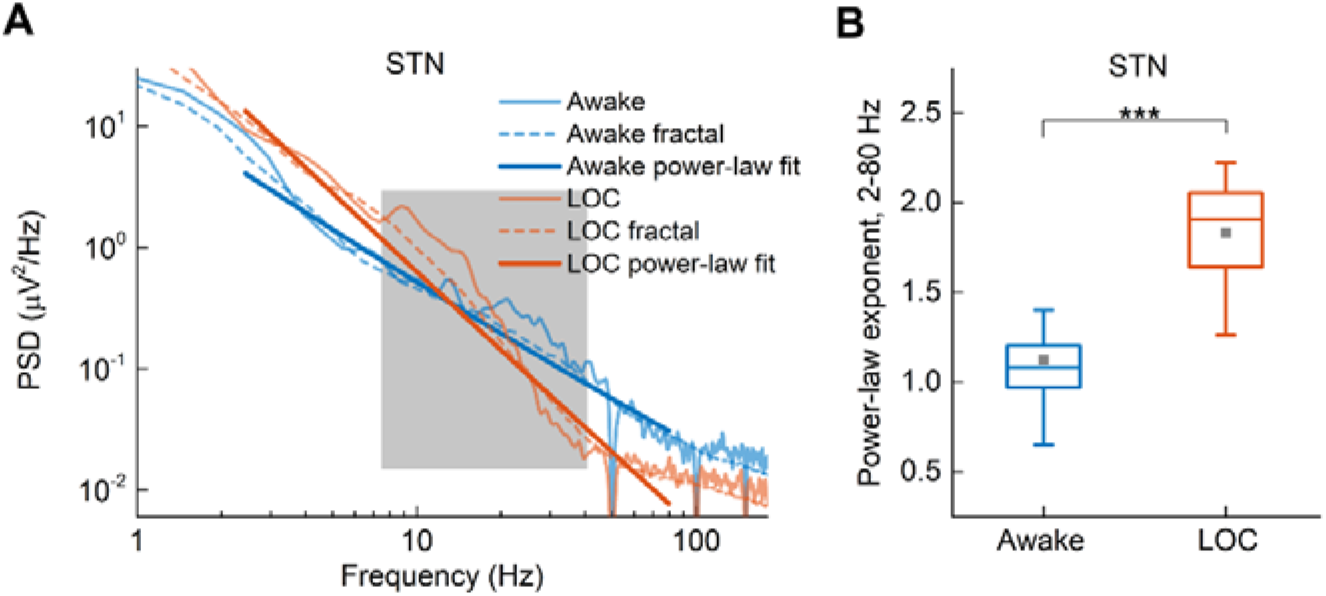
Increased power-law exponent at 2-80 Hz in STN LFPs at LOC state compared with awake state. (A) Representative power spectra, corresponding fractal components and power-law fit of the STN LFPs in awake and LOC states. The range with local power peaks was marked by the shadow. (B) Boxplot showing power-law exponent in STN LFPs across subjects in awake and LOC states. ***, *p*<0.001.

### Changes of neural activity in the STN induced by intubation in anaesthesia

Intubation is an arousal intervention in anaesthesia. Figure 3A shows representative spectrograms in the STN LFP and frontal EEG activity around intubation procedure under anaesthesia showing apparent increased beta activity in the STN at intubation. Group-level spectral analysis (n = 20, ten subjects) revealed significant increase in beta and broad band high-gamma power in the STN (low beta: *p* = 0.0257, high-beta: *p* = 0.0064, high-gamma: *p* = 0.033; FDR correction). In comparison, we did not observe any significant changes over the frontal cortex comparing intubation against the status of loss of consciousness (Fig. 3B and C). The imaginary coherence between STN and frontal cortex showed slight but non-significant decrease in alpha band in intubation compared with LOC state (Fig. 3D).

**Figure 3.**
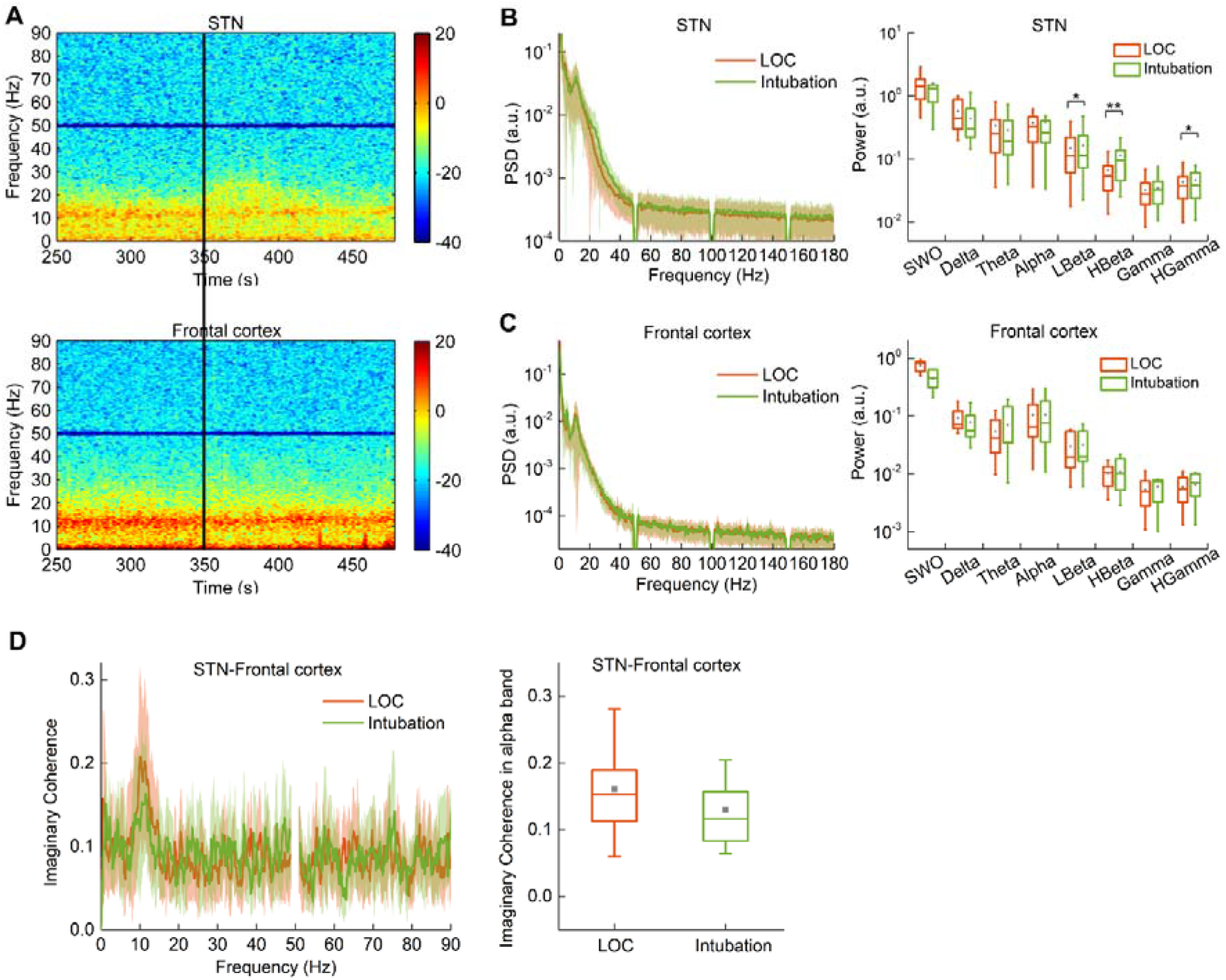
Intubation was associated with increased activity in the beta frequency and broadband high-gamma band in the STN, but not over the frontal cortex. (A) Representative time-frequency spectrograms of STN LFPs and frontal cortical EEG around intubation in anaesthesia. The vertical black line marks the time of starting intubation. (B) Group spectra and boxplot showing power in different frequency bands of STN LFPs across subjects in LOC state. (C) Group spectra and boxplot showing power in different frequency bands of STN LFPs across subjects in intubation state. (D) Group imaginary coherence between STN and frontal cortex across subjects in awake and LOC states. *, *p*<0.05; **, *p*<0.01.

### Changes in power-law exponent in STN LFPs and frontal EEG induced by intubation

Representative power spectra of the STN LFPs and frontal EEG and their linear fitting in log-log coordinates at LOC and intubation states were shown in Figures 4A and C. Compared to LOC state, averaged power-law exponent significantly decreased from 1.83±0.30 (range 1.26-2.21) to 1.66±0.30 (range 1.09-1.97) at intubation state in the STN (*p* = 0.0012; FDR correction) (Fig. 4B). However, the power-law exponent is still higher than that during awake (*p* < 0.0001; FDR correction) (Fig. 4B), suggesting scaled changes in the power-law exponent of STN LFPs associated with different levels of consciousness and arousal during anaesthesia. However, the effect of intubation on power-law exponent in frontal cortical EEG was non-significant (Fig. 4D).

**Figure 4.**
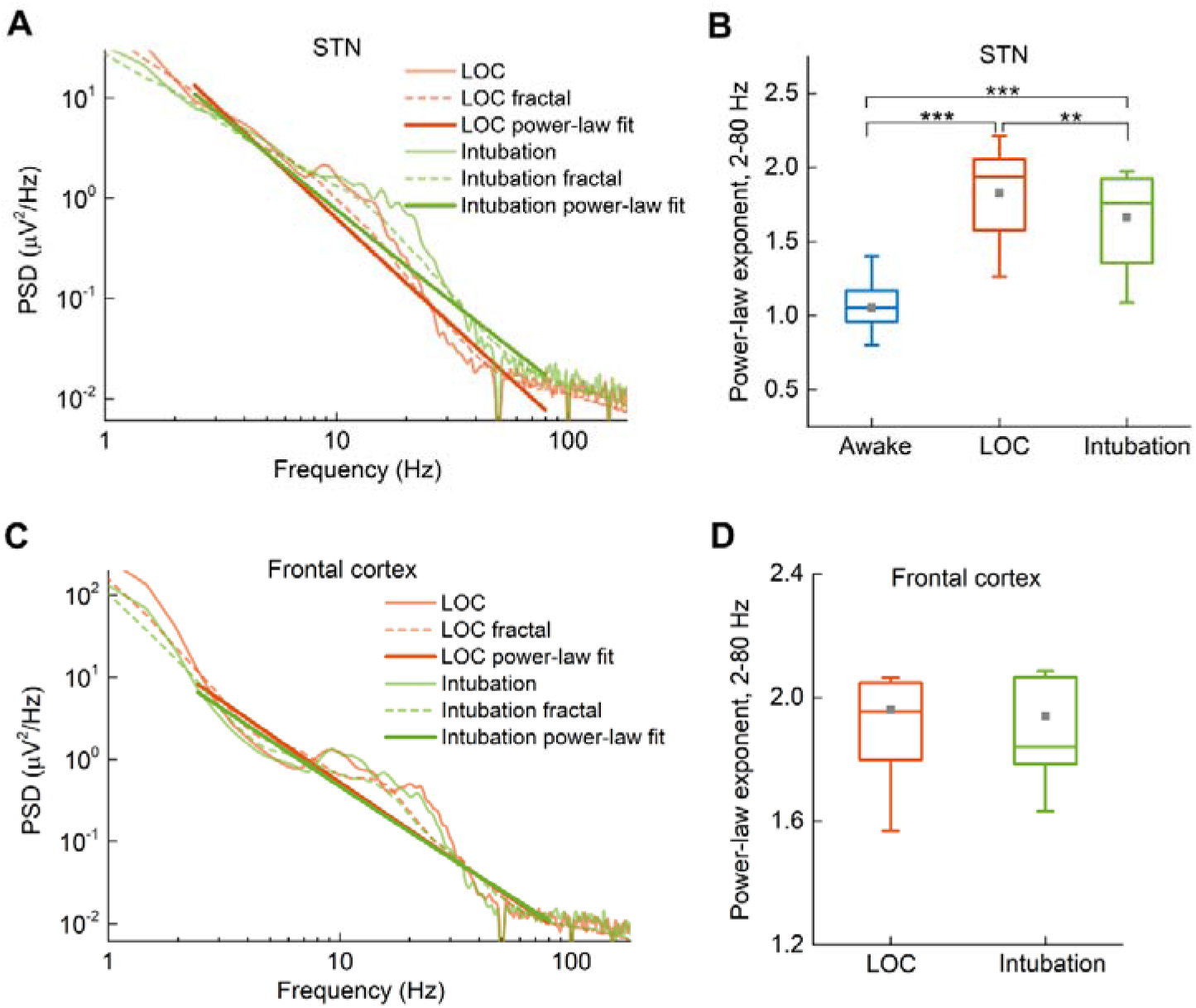
Power-law difference in STN LFPs and frontal EEG between LOC and intubation states. (A) Representative power spectra, corresponding fractal components and power-law fit of the STN LFPs in LOC and intubation states. (B) Boxplot showing power-law exponent in STN LFPs across subjects in awake, LOC and intubation states. (C) Representative power spectra, corresponding fractal components and power-law fit of the frontal EEG in LOC and intubation states. (D) Boxplot showing power-law exponent in frontal EEG across subjects in LOC and intubation states. **, *p*<0.01; ***, *p*<0.001.

### Dynamic power-law fluctuation of STN neural activity during general anaesthesia

Computational modelling and recent experimental data suggested that the power-law exponent of the local field potentials indicate excitation/inhibition balance in the neural circuits ^29^. We therefore further hypothesized that the dynamic fluctuation of the power-law exponent in the STN will be correlated with high-gamma activity which reflects the local population firing and multi-unit activity ^35–38^ during general anaesthesia. To test this hypothesis, we applied a sliding time window with a length of 2 s and overlap of 1 s and quantified the power-law exponent in the 2-80 Hz and high-gamma power (80-180 Hz) for each time window. We found that power-law exponent gradually increased and then plateaued at about 70s after the loss of consciousness; the power-law exponent reduced rapidly following intubation in anaesthesia but remained higher than during awake (Fig. 5A). In addition, the power at high-gamma frequency band varied in a coherent way that followed the power-law relationship. We estimated the within-subject correlation between power-law exponent and high-gamma power in anaesthesia (100 s before intubation and 100 s after intubation). This analysis identified significant correlation in 15 out of 20 STNs, with average correlation coefficient of −0.25 ± 0.14 (Mean ± SE across all STNs) (Fig. 5B and C). In addition, there is significant correlation between the change in the high frequency power and the change in the lower frequency power-law exponent induced by intubation relative to LOC state across subjects (Fig. 5D).

**Figure 5.**
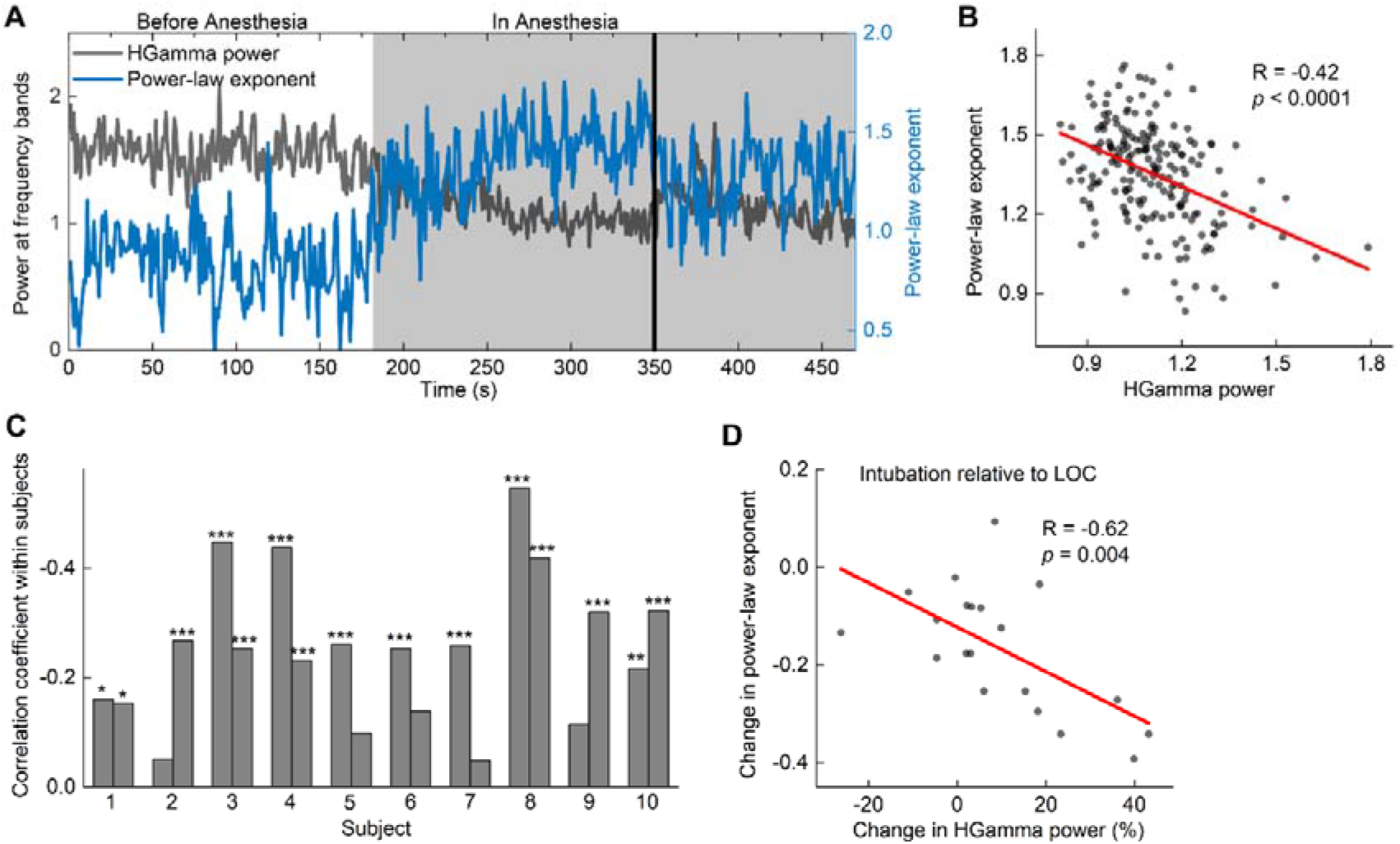
Dynamic power-law exponent and broadband high-gamma power tracking the change of E/I balance in the STN. (A) Representative co-fluctuations of power-law exponent and high-gamma power in a typical STN LFPs during induction of anaesthesia and intubation. The vertical black line marks the time of starting intubation. (B) The scatterplot showing correlation between power-law exponent and high-gamma power in anaesthesia shown in (A). (C) Correlations between power-law exponent and high-gamma power for subjects. (D) The correlation between the change in high-gamma power and the change in power-law exponent in STN LFPs at intubation relative to LOC state across subjects. *, *p*<0.05; **, *p*<0.01; ***, *p*<0.001.

## Discussion

In the present study, we used the unique opportunity afforded by synchronous electrophysiological recordings directly from human STN and frontal cortex to characterize changes in both oscillatory and arrhythmic activities in basal ganglia and cortical dynamics associated with consciousness in general anaesthesia. We showed that, (1) unconsciousness state induced by propofol induced changes in multiple frequency bands in human STN LFP. These changes include increased oscillations in low frequencies including SWO, delta, theta, and alpha bands, and reduced oscillation in beta frequency band, as well as reduced broadband high-frequency activity. There is also increased coherence between STN and frontal cortex, but only in the alpha band. (2) LFP power-law exponent in the STN between 2 and 80 Hz significantly increased at loss of consciousness compared to awake state and temporally decreased following intubation, but it was still higher than the awake state. The dynamic changes in the power law exponent correlated with fluctuation of broadband high-frequency activity. Our finding coincides with previous computational modelling and animal studies showing increased frontal corticothalamic alpha coherence associated with consciousness ^13, 39^. Our results are also consistent with the previous study showed reduced beta band activity in STN during general anaesthesia ^19^. Furthermore, our results highlighted more comprehensive changes in STN during general anaesthesia and intubation, which was not observed in the cortical EEG, suggesting a more complicated role of STN in the modulation of consciousness.

Our study is novel in three ways. First, we revealed consistent variation of power-law distribution in the 2-80 Hz range in the PSD of STN LFPs in general anaesthesia. Most previous studies on the EEG, ECoG and MEG recordings have focused on the narrow band oscillatory activities, treating the scale-free activities following the power law as recording noise. The power-law scaling of the PSD could be explained by different mechanisms according to different frequency ranges ^22, 25^. Recent evidence from computational modelling and in vivo recordings suggested that the power-law exponent in low frequency bands reflects the balance between excitation and inhibition ^29^. The balance between excitation and inhibition plays a crucial role in the brain functioning of neuronal networks ^40^, and the power-law exponent in the cortical LFPs have been shown to be modulated in anaesthesia ^29, 30, 41^. This is the first study, to our knowledge, suggests that the power-law exponent in STN LFPs is consistently modulated by anaesthetic-induced unconsciousness in humans. This is also consistent with the hypothesis that the power-law exponent reflects excitation/inhibition balance. Propofol may have decreased excitation/inhibition ratio in the STN in anaesthesia, which is reflected in the steeper power-law slope in the STN LFPs.

Second, we identified reduced broadband high-gamma activity in the STN in anaesthesia. Broadband high-frequency activity without a narrow power peak in the spectrum is considered a reflection of arrhythmic activity ^25^, and has been suggested to potentially reflect the local population firing activity ^42^ or with neuronal spiking rate ^35, 36^. Thus, the attenuation of high-gamma power by propofol we observed in the STN in this study likely reflects reduced neuronal firing rates in anaesthesia, which has been reported in recent studies using microelectrode recordings (MER) in the STN ^43, 44^.

In addition, we show that the dynamic temporal fluctuations in power law exponent in the lower frequency band correlated with the changes in broad-band high frequency power, again supporting the hypothesis that anaesthesia can lead to changes in the excitation/inhibition balance in STN which may in turn reduce nucleus output in the form of multi-unit activity. On the other hand, there was no correlation between high frequency power and power of low frequency activities (alpha band) during anaesthesia (not shown) which was suggested to be a neural correlate of synchronized afferent input to the nucleus ^35, 45^.

Third, we also investigated the changes of neural signatures in STN at intubation state during general anaesthesia. This is valuable, since clinically relevant noxious stimuli such as tracheal intubation or skin incision may induce intraoperative awareness ^46–48^. The efficacy of different approaches to detect and prevent intraoperative awareness with recall, including the use of EEG-based monitoring such as bispectral index (BIS), has been controversial in several prospective randomized controlled trials ^49–52^. The frontal EEG biomarkers also failed to detect awareness following intubation ^46, 47, 53^. Our findings from STN LFPs demonstrated that there were neural responses in the subcortical areas to intubation in anaesthesia, potentially advancing our understanding of intraoperative awareness in the future. There are some limitations in this study. First, our experiments were conducted within clinical settings in which anaesthetic management was administered according to the anaesthetist’s discretion, rather than in a controlled, titrated manner. This prevented us from correlating the neural signatures with precise consciousness level. We mitigated these defects by examining the approximately steady states prior to and after induction of anaesthesia. Moreover, we used intubation state as an arousing intervention to investigate whether there was a reversal of the patterns, partially supporting the findings of biomarkers for consciousness, and the dynamic changes over time of these neural signatures were achieved by combining these measures and a time-windowing approach. Future studies would benefit from target-controlled infusion approach during induction of anaesthesia to have a detailed investigation of concentration-dependent effects featuring hierarchic states of consciousness. Second, the electrophysiological recordings were from PD patients in this study, raising a consideration of whether the reported changes in the STN were related to PD symptoms. However, the patients were on medication for controlling PD symptoms during recording, partly mitigating the effect of disease. And effects of propofol on STN activity were highly consistent with those observed in cortical areas. This suggests that the reported characteristics here is likely to partly generalize to the healthy brain. It should also be acknowledged that the power-law exponent quantified in this study is not independent from the power of different sub-frequency bands, including alpha, beta and mid-gamma bands. However, the power-law exponent takes into account the relationship between the power of different frequency band, and can provide an estimate of the high gamma activity.

In conclusion, we characterized STN neural activity from two profiles, including oscillatory features and non-oscillatory (scale-free) features during anaesthesia. These findings provide important electrophysiological evidence in humans of the neural mechanisms regarding basal ganglia for propofol-induced unconsciousness.

## Author Contributions

Study design: Y.H., K.H., H.T.; Data collection: Y.H., K.H., A.L.G., X.M., M.J.G., J.J.F., Y.P., S.M., P.H., S.Z., D.L., T.Z.A., B.S.; Data analysis: Y.H., H.T.; Data interpretation: all authors; Drafting of the manuscript: Y.H., H.T.; Revising of the manuscript: all authors.

## Acknowledgements

We acknowledge Dr. Simon Yarrow for conducting anaesthesia on some of the patients, Prof. Peter Brown for constructive and critical comments.

## Declaration of Interests

The authors report no competing interests

## Funding

This work was supported by Shanghai Pujiang Program (K.H, 19PJ1407500), Medical and Engineering Cross Research Fund from Shanghai Jiao Tong University (K.H, YG2019QNA31), and The National Institute for Health Research (NIHR) Oxford Biomedical Research Centre. B.S. is funded by National Natural Science Foundation of China (81471482), Shanghai Jiao Tong University School of Medicine – Institute of Neuroscience (SHSMU-ION) Research Center for Brain Disorders. M.J.G. is funded by the Academy of Medical Sciences, Starter Grant for Clinical lecturers.

